# Neurophysiological signatures of cortical micro-architecture

**DOI:** 10.1101/2023.01.23.525101

**Authors:** Golia Shafiei, Ben D. Fulcher, Bradley Voytek, Theodore D. Satterthwaite, Sylvain Baillet, Bratislav Misic

## Abstract

Systematic spatial variation in micro-architecture is observed across the cortex. These micro-architectural gradients are reflected in neural activity, which can be captured by neurophysiological time-series. How spontaneous neurophysiological dynamics are organized across the cortex and how they arise from heterogeneous cortical micro-architecture remains unknown. Here we extensively profile regional neurophysiological dynamics across the human brain by estimating over 6 800 timeseries features from the resting state magnetoencephalography (MEG) signal. We then map regional time-series profiles to a comprehensive multi-modal, multi-scale atlas of cortical micro-architecture, including microstructure, metabolism, neurotransmitter receptors, cell types and laminar differentiation. We find that the dominant axis of neurophysiological dynamics reflects characteristics of power spectrum density and linear correlation structure of the signal, emphasizing the importance of conventional features of electromagnetic dynamics while identifying additional informative features that have traditionally received less attention. Moreover, spatial variation in neurophysiological dynamics is colocalized with multiple micro-architectural features, including genomic gradients, intracortical myelin, neurotransmitter receptors and transporters, and oxygen and glucose metabolism. Collectively, this work opens new avenues for studying the anatomical basis of neural activity.

## INTRODUCTION

Signals, in the form of electrical impulses, are perpetually generated, propagated and integrated via multiple types of neurons and neuronal populations [1, 2]. The wiring of the brain guides the propagation of signals through networks of nested polyfunctional neural circuits [3, 4]. The resulting fluctuations in membrane potentials and firing rates ultimately manifest as patterned neurophysiological activity [5–7].

A rich literature demonstrates links between cortical micro-architecture and dynamics. Numerous studies have investigated the cellular and laminar origins of cortical rhythms [8–13]. For instance, electro- and magneto-encephalography (EEG/MEG) signals appear to be more sensitive to dipoles originating from pyramidal cells of cortical layers II-III and V [14, 15]. Moreover, specific time-series features of neuronal electrophysiology depend on neuron type, morphology and local gene transcription, particularly genes associated with ion channel regulation [16–18]. However, previous studies have mostly focused on single or small sets of features-of-interest, often mapping single micro-architectural features to single dynamical features. Starting with the discovery of 8-12 Hz alpha rhythm in the electroencephalogram [19], conventional time-series analysis in neuroscience has typically focused on canonical electrophysiological rhythms [20–24]. More recently, there has also been a growing interest in studying the intrinsic timescales that display a hierarchy of temporal processing from fast fluctuating activity in unimodal cortex to slower encoding of contextual information in trans-modal cortex [25–34]. How ongoing neurophysiological dynamics arise from specific features of neural circuit micro-architecture remains a key question in neuroscience [1, 2, 12].

Recent analytic advances have opened new opportunities to perform neurophysiological time-series phenotyping by computing comprehensive feature sets that go beyond power spectral measures, including measures of signal amplitude distribution, entropy, fractal scaling and autocorrelation [35–40]. Concomitant advances in imaging technologies and data sharing offer new ways to measure brain structure with unprecedented detail and depth [41–43], including gene expression [44], myelination [45, 46], neurotransmitter receptors [47–54], cytoarchitecture [55–57], laminar differentiation [56, 58], cell type composition [44, 59, 60], metabolism [61, 62] and evolutionary expansion [63, 64].

Here we comprehensively characterize the dynamical signature of neurophysiological activity and relate it to the underlying micro-architecture by integrating multiple, multimodal maps of human cortex. We first derive cortical spontaneous neurophysiological activity using source-resolved magnetoencephalography (MEG) from the Human Connectome Project (HCP; [65]). We then apply highly comparative time-series analysis (`hctsa`; [35, 36]) to estimate a comprehensive set of time-series features for each brain region (Fig. 1). At the same time, we construct a micro-architectural atlas of the cortex that includes maps of microstructure, metabolism, neurotransmitter receptors and transporters, laminar differentiation and cell types (Fig. 2). Finally, we map these extensive micro-architectural and dynamical atlases to one another using multivariate statistical analysis.

**Figure 1.**
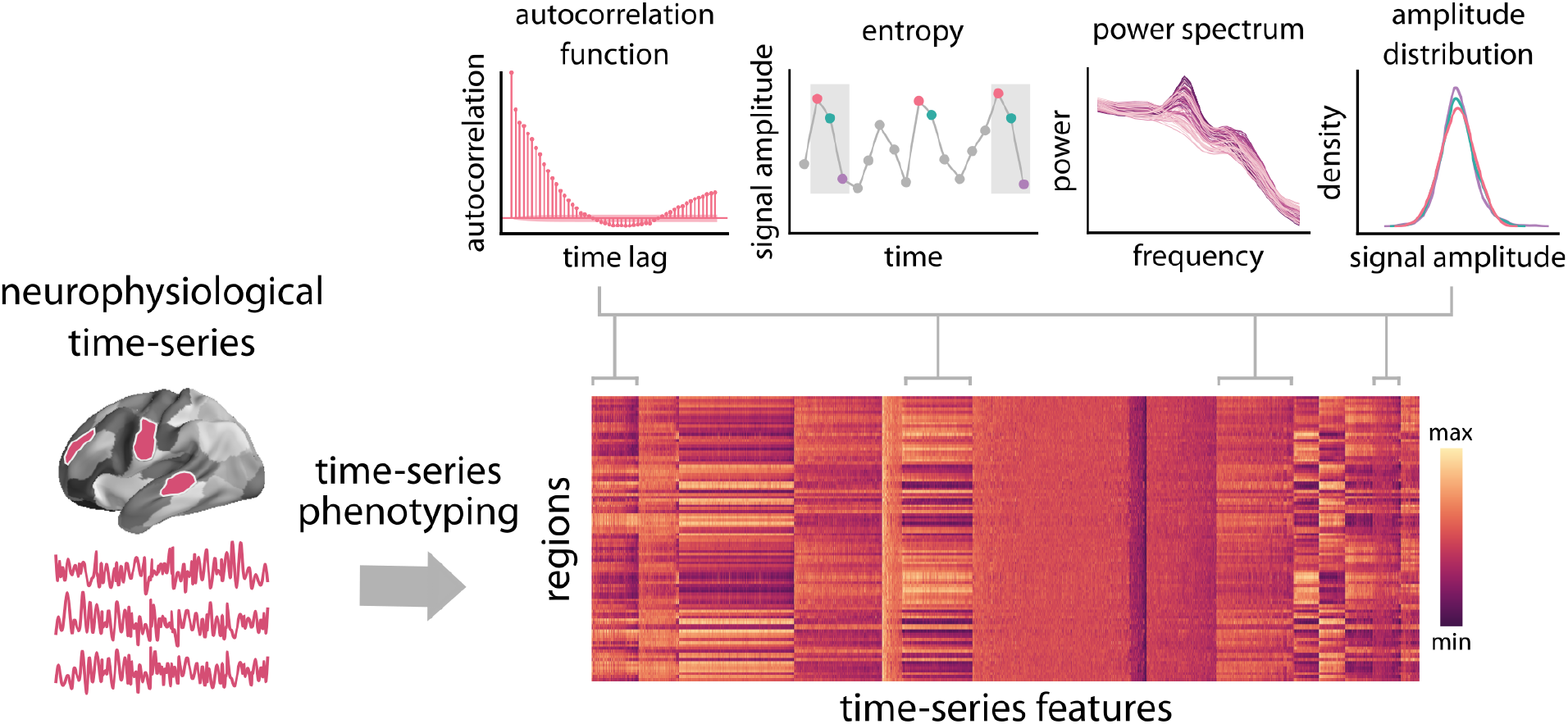
Feature-based representation of neurophysiological time-series. Highly comparative time-series analysis (`hctsa`; [35]) toolbox was used to perform time-series feature extraction on regional MEG time-series. This time-series phenotyping procedure generated 6 880 time-series features for each region, including measures of autocorrelation, entropy, power spectrum and amplitude distribution.

**Figure 2.**
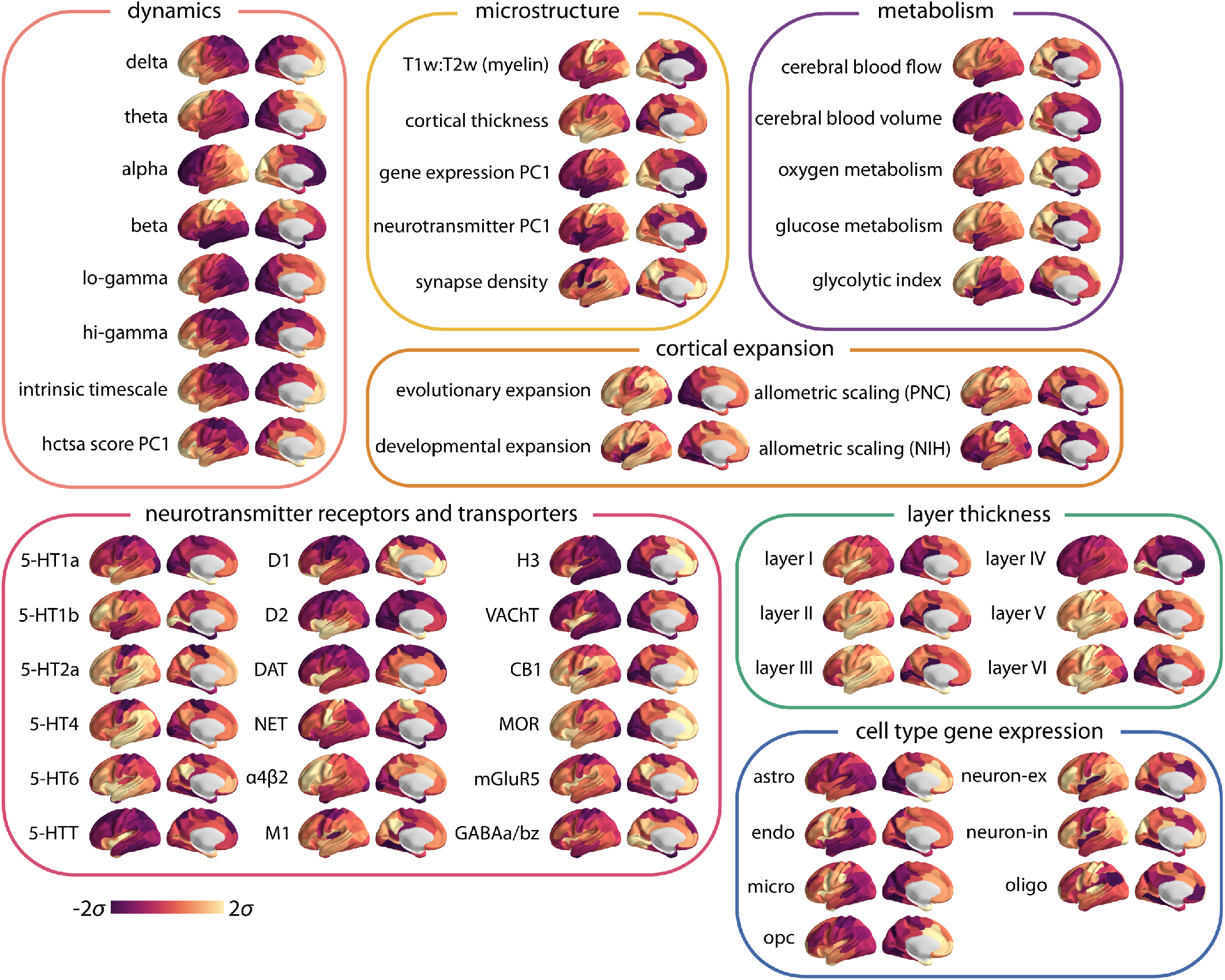
Multimodal brain maps. neuromaps toolbox [43], BigBrainWarp toolbox [57], the Allen Human Brain Atlas (AHBA [44]) and the abagen toolbox [68] were used to compile a set of 45 micro-architectural brain maps, including measures of microstructure, metabolism, cortical expansion, receptors and transporters, layer thickness and cell type-specific gene expression (astro = astrocytes; endo = endothelial cells; micro = microglia; neuron-ex = excitatory neurons; neuron-in = inhibitory neurons; oligo = oligodendrocytes; and opc = oligodendrocyte precursors) (see *Methods* for more details). Note that the microstructure maps include principal gradients of gene expression and neurotransmitter profiles, for which we have also separately included feature sub-sets (specific receptor maps and cell type specific gene expression). Brainstorm software was used to pre-process the resting-state MEG data and obtain power maps at six canonical electrophysiological bands (i.e., delta (*δ*: 2-4 Hz), theta (*θ*: 5-7 Hz), alpha (*α*: 8-12 Hz), beta (*β*: 15-29 Hz), low gamma (lo-*γ*: 30-59 Hz), and high gamma (hi-*γ*: 60-90Hz)) [66] (see *Methods* for more details). FOOOF algorithm was used to parametrize power spectral density and estimate the intrinsic timescale [21, 30] (see *Methods* for more details). Note that log-10 transformed intrinsic timescale map is shown here. Principal component analysis was used to estimate the principal component of the neurophysiological time-series features obtained from the `hctsa` toolbox (see Fig. 3). All obtained brains maps are depicted across the cortex at 95% confidence interval (Schaefer-100 atlas [67]).

## RESULTS

Regional neurophysiological time-series were estimated by applying linearly constrained minimum variance (LCMV) beamforming to resting state MEG data from the Human Connectome Project (HCP; [65]) using Brainstorm software [66](see *Methods* for details). Highly comparative time-series analysis (`hctsa`; [35, 36]) was then applied to regional time-series to estimate 6 880 time-series features for 100 cortical regions from the Schaefer-100 atlas [67]. The list of time-series features includes, but is not limited to, statistics derived from the autocorrelation function, power spectrum, amplitude distribution, and entropy estimates (Fig. 1). This time-series phenotyping procedure yields a comprehensive, data-driven “fingerprint” of regional dynamical properties of each brain region.

To estimate a comprehensive set of multimodal micro-architectural features, we used the recently-developed neuromaps toolbox [43] as well as the BigBrainWarp toolbox [57], the Allen Human Brain Atlas (AHBA [44]) and the abagen toolbox [68] to transform and compile a set of 45 features, including measures of microstructure, metabolism, cortical expansion, receptors and transporters, layer thickness and cell type-specific gene expression (Fig. 2).

In subsequent analyses, we first assess the topographic organization of neurophysiological dynamics by quantifying the dominant patterns of variations in resting-state MEG time-series properties. We then characterize the signature of neurophysiological dynamics with respect to micro-architectural attributes across the cortex. Finally, we perform sensitivity analyses to investigate potential effects of confounding factors on the findings, such as signal-to-noise ratio and parcellation resolution (see *Sensitivity analysis* for details).

### Topographic distribution of neurophysiological dynamics

The `hctsa` time-series phenotyping procedure generated 6 880 time-series features per brain region. Since the identified time-series features potentially capture related dynamical behaviour and contain groups of correlated properties, we first sought to identify dominant macroscopic patterns or gradients of neurophysiological dynamics [38]. Applying principal component analysis (PCA) to the group-average region × feature matrix, we find evidence of a single dominant component that captures 48.7%of the variance in regional time-series features (Fig. 3a). The dominant component or “gradient” of neurophysiological dynamics (PC1) mainly spans the posterior parietal cortex and sensory-motor cortices on one end and the anterior temporal, orbitofrontal and ventromedial cortices on the other end (Fig. 3a). Focusing on intrinsic functional networks, we find that the topographic organization of the dominant neurophysiological dynamics varies along a sensory-fugal axis from dorsal attention, somatomotor and visual networks to limbic and default mode networks [69] (Fig. 3a).

**Figure 3.**
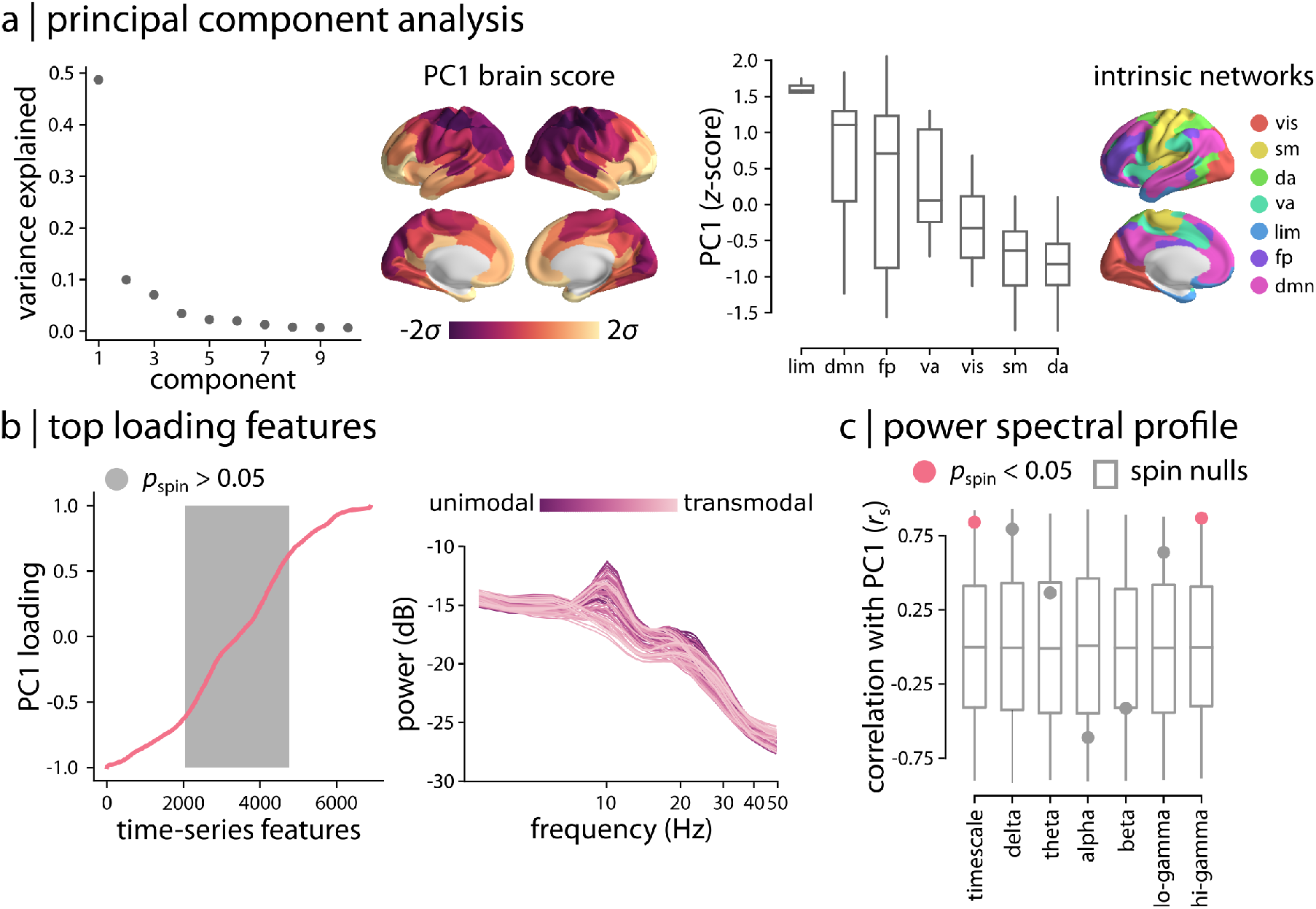
Topographic distribution of neurophysiological dynamics. (a) Principal component analysis (PCA) was used to identify linear combinations of MEG time-series features with maximum variance across the cortex. The fist principal component (PC1) accounts for 48.7% of the total variance in neurophysiological time-series features. The spatial organization of the dominant time-series features captured by PC1 is depicted across the cortex at 95% confidence interval. The distribution of PC1 brain score is also depicted for intrinsic functional networks [69]. (b) To examine the feature composition of the time-series features captured by PC1, feature loadings were estimated as the correlation coefficients between each `hctsa` time-series feature and PC1 brain score. PC1 loadings are depicted for the time-series features (ordered by their individual loadings). Grey background indicates non-significant features based on 10 000 spatial autocorrelation-preserving permutation tests (i.e., “spin tests” [70, 71]; FDR-corrected). The top loading features were mainly related to the power spectrum of regional time-series and its structure. Regional power spectral densities are depicted, with each line representing a brain region. Regions are coloured by their position in the putative unimodal–transmodal hierarchy [72]. (c) To contextualize the principal component of variation in MEG timeseries features, PC1 brain score was correlated with MEG power maps at 6 canonical frequency bands and intrinsic timescale. The observed correlations are shown by filled circles and are compared to their corresponding null distributions of correlations obtained from 10 000 spatial autocorrelation-preserving permutation tests (“spin nulls” depicted as grey box plots). PC1 score is significantly correlated with intrinsic timescale and hi-gamma power (FDR-corrected). *r*_s_ denotes the Spearman’s rank correlation coefficient. Intrinsic networks: vis = visual; sm = somatomotor; da = dorsal attention; va = ventral attention; lim = limbic; fp = frontoparietal; dmn = default mode.

We next investigated the top-loading time-series features on the first component, using the univariate correlations between each of the original feature maps and the PC1 map (i.e., PCA loadings). All correlations were statistically assessed using spatial autocorrelationpreserving null models (“spin tests” [70, 71]; see *Methods* for details). Fig. 3b shows that numerous features are positively and negatively correlated with PC1; the full list of features, their correlation coefficients and *p*-values are available in the online Supplementary File S1. Inspection of the top loading features reveals that the majority are statistics derived from the structure of the power spectrum or closely related measures. Examples include power in different frequency bands, parameters of various model fits to the power spectrum, and related measures, such as the shape of the autocorrelation function and measures of fluctuation analysis. Fig. 3b shows how the power spectrum varies across the cortex, with each line representing a brain region. Regions are coloured by their position in the putative unimodal–transmodal hierarchy [72]; the variation visually suggests that uni-modal regions display more prominent alpha (8-12 Hz) and beta (15-29 Hz) power peaks. Collectively, these results demonstrate that the traditional focus of electrophysiological time-series analysis on statistics of the power spectrum is consistent with the dominant variations in MEG dynamics captured by the diverse library of `hctsa` time-series features.

Given the hierarchical organization of PC1 and its close relationship with power spectral features, we directly tested the link between PC1 and conventional band-limited power spectral measures [21–23], as well as intrinsic timescale [30]. Fig. 3c shows the correlations between PC1 and delta (2-4 Hz), theta (5-7 Hz), alpha (8-12 Hz), beta (15-29 Hz), lo-gamma (30-59 Hz) and hi-gamma (60-90Hz) power maps, and intrinsic neural timescale [28, 30–34, 73]. We find that PC1 has high spatial correlations with most of those maps (|*r*| > 0.36), and significant correlations with intrinsic timescale (*r*_s_ = 0.84, *p*_spin_ = 0.03; FDR-corrected) and hi-gamma (*r*_s_ = 0.87, *p*_spin_ = 0.005; FDR-corrected). The results were consistent when we used band-limited power maps that were adjusted for the aperiodic component of the power spectrum as opposed to the total power [21] (Fig. S1). The fact that PC1 correlates with intrinsic timescale is consistent with the notion that both capture broad variations in the power spectrum. Note that we focused on PC1 because the other components (PC2 and above) accounted for 10% or less of the variance in time-series features and they were not significantly associated with `hctsa` time-series features.

### Neurophysiological signatures of micro-architecture

How do these regional neurophysiological time-series features map onto multimodal micro-architectural features? To address this question, we implemented a multivariate partial least squares analysis (PLS; [74, 75]) that integrates multiple multimodal brain maps into the analysis and seeks to identify linear combinations of time-series features and linear combinations of micro-architectural features that optimally covary with one another. Fig. 4a shows that the analysis identifies multiple such combinations, termed latent variables. Statistical significance of each latent variable was assessed using spatial autocorrelation-preserving permutation tests [71, 76]. The first latent variable was statistically significant, capturing the greatest covariance between timeseries and micro-architectural features (covariance explained = 75.4%, *p*_spin_ = 0.011).

**Figure 4.**
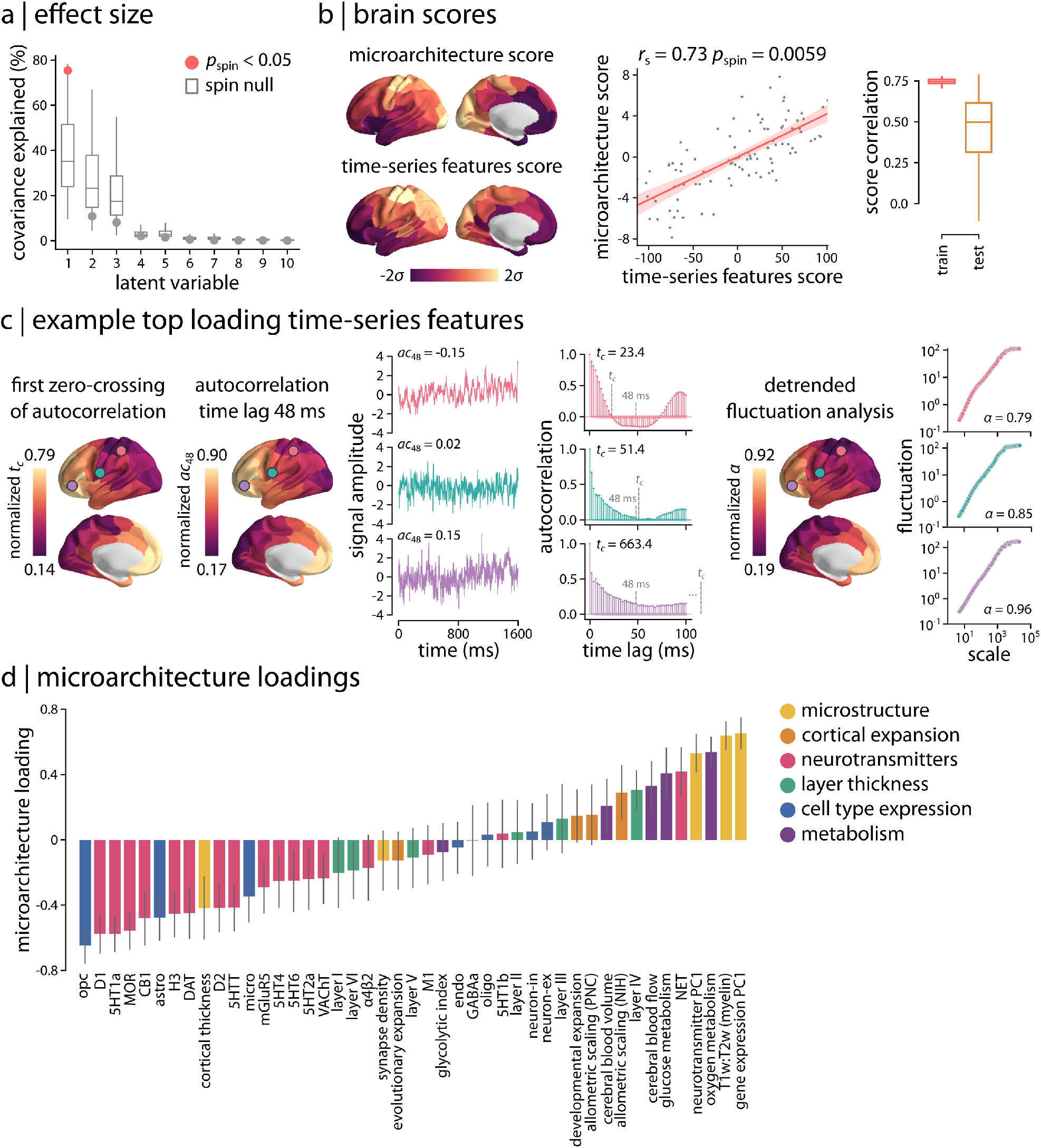
Neurophysiological signature of micro-architecture. (a) Partial least square (PLS) analysis was used to assess the multivariate relationship between micro-architectural and time-series features. PLS identified a single significant latent variable (*p*_spin_ = 0.011, covariance explained = 75.4%). (b) Spatial patterns of micro-architecture and time-series features scores are depicted for the first latent variable. The two brain score maps are significantly correlated. To assess the out-of-sample correlation of brain scores, a distance-dependent cross-validation analysis was used (see *Methods*). Micro-architecture and time-series feature scores are consistently correlated in both training set (75% of regions; mean *r*_s_ = 0.75) and test set (25% of regions; mean r*_s_* = 0.5). *r_s_* denotes the Spearman’s rank correlation coefficient; linear regression line is added to the scatter plot for visualization purposes only. (c) Top loading time-series features were mainly related to measures of self-correlation or predictability of the signal. Three examples of top loading features are depicted across the cortex. Left: first zero-crossing time point of the autocorrelation function, *t_c_*, and linear autocorrelation at a lag of 48 ms, *ac*_48_; right: the scaling exponent of detrended fluctuation analysis, *α*. Short segments of raw time-series, autocorrelation functions, and fluctuation analysis plots (log-log plot of detrended fluctuations at multiple timescales) are also shown for a randomly selected participant at three cortical regions (circles on the brain surface: pink ≈ 5^*th*^ percentile, green ≈ 50^*th*^ percentile, purple ≈ 95^*th*^ percentile). Time points corresponding to zero-crossing point (*t_c_*) and 48 ms are indicated with grey dashed lines on the autocorrelation function plots. (d) Micro-architectural feature loadings are shown for each set of brain maps.

Fig. 4b shows the spatial topography of time-series features and micro-architectural scores for the first latent variable. These are the weighted sums of the original input features according to the weighting identified by the latent variable. The correlation between the score maps is maximized by the analysis (*r*_s_ = 0.73, *p*_spin_ = 0.0059). We therefore sought to estimate whether the same mapping between time-series and micro-architectural features can be observed out-of-sample. We adopted a distance-dependent cross-validation procedure where “seed” regions were randomly chosen and the 75% most physically proximal regions were selected as the training set, while the remaining 25% most physically distal regions were selected as the test set [76] (see *Methods* for more details). For each train-test split, we fit a PLS model to the train set and project the test set onto the weights (i.e., singular vectors) derived from the train set. The resulting test set scores are then correlated to estimate an out-of-sample correlation coefficient. Fig. 4b shows that micro-architecture and time-series feature scores are correlated in training set (mean *r*_s_ = 0.75) and test set (mean *r*_s_ = 0.5), demonstrating consistent findings in out-of-sample analysis.

We next examined the corresponding time-series and micro-architecture feature loadings and identified the most contributing features to the spatial patterns captured by the first latent variable (Fig. 4c,d). The top loading time-series features were mainly related to measures of self-correlation or predictability of the MEG signal. The self-correlation measures mostly reflect the linear correlation structure of neurophysiological timeseries, particularly long-lag autocorrelations (at lags > 15 time steps, or > 30 ms). A wide range of other highly weighted time-series features captured other aspects of signal predictability, including measures of the shape of the autocorrelation function (e.g., the time lag at which the autocorrelation function crosses zero), how the autocorrelation structure changes after applying simple local forecasting models (e.g., residuals from fitting linear models to rolling 5-time-step, or 10 ms, windows), scaling properties assessed using fluctuation analysis (e.g., scaling of signal variance across timescales), and measures derived from a wavelet decomposition (e.g., wavelet coefficients at different timescales). The full list of time-series feature loadings for the first latent variable is available in the online Supplementary File S2.

To illustrate the spatial distribution of highly contributing time-series features, Fig. 4c shows three top-loading features that mirror the spatial variation of the first PLS latent variable. For example, Fig. 4c,*left* depicts the distribution of the group-average first zero-crossing point of the autocorrelation function. The autocorrelation function of the unimodal cortex (marked with a pink circle) crosses zero autocorrelation at a lower lag than the transmodal cortex (marked with a purple circle), suggesting faster autocorrelation decay and longer correlation length in transmodal cortex than in unimodal cortex. Another example is the linear autocorrelation of the MEG signal at longer time lags. Fig. 4c,*left* shows autocorrelation at a lag of 48 ms (24 time steps), demonstrating lower autocorrelation in unimodal cortex and higher autocorrelation in transmodal cortex. Finally, we examined the scaling exponent, *α*, estimated using detrended fluctuation analysis as the slope of a linear fit to the log-log plot of the fluctuations of the detrended signal across timescales [77, 78]. Fig.4c,*right* depicts this scaling exponent across the cortex, which exhibits a similar spatial pattern as the previous two examples, indicating lower self-correlation in unimodal cortex (pink circle) compared to transmodal cortex (purple circle). Other variations of fluctuation analysis also featured heavily in the list of top-loading features, including goodness of fit of the linear fit, fitting of multiple scaling regimes, and different types of detrending and mathematical formulation of fluctuation size.

Fig. 4d shows the corresponding micro-architectural loadings. The most contributing micro-architectural features to the spatial patterns captured by the first latent variable are the principal component of gene expression (gene expression PC1; a potential proxy for cell type distribution [44, 76, 79]), T1w/T2w ratio (a proxy for intracortical myelin), principal component of neurotransmitter receptors and transporters (neurotransmitter PC1), and oxygen and glucose metabolism (strong positive loadings). We also find high contributions (strong negative loadings) from specific neurotransmitter receptor and transporters, in particular metabotropic serotonergic and dopaminergic receptors, as well as from cell type-specific gene expression of oligodendrocyte precursors (opc), which are involved in myelinogenesis [80–84]. Consistent findings were obtained when we used univariate analysis to relate regional time-series features and the top loading micro-architectural maps, in particular principal component of gene expression and T1w/T2w ratio, which have previously been extensively studied as archetypical micro-architectural gradients [30, 41, 79, 85, 86] (Fig. S2). Altogether, this analysis provides a comprehensive chart or “lookup table” of how micro-architectural and time-series feature maps are associated with one another. These results demonstrate that cortical variation in multiple micro-architectural attributes manifests as a gradient of time-series properties of neurophysiological activity, particularly the properties that reflect the long-range self-correlation structure of the signal.

### Sensitivity analysis

To assess the extent to which the results are affected by potential confounding factors and methodological choices, we repeated the analyses using alternative approaches. First, to ensure that the findings are not influenced by MEG signal-to-noise ratio (SNR), we calculated SNR at each source location using a noise model that estimates how sensitive the source-level MEG signal is to source location and orientation [87, 88]. We performed two follow-up analyses using the SNR map (Fig. S3): (1) SNR was first compared with the full set of MEG time-series features using mass univariate Pearson correlations. Time-series features that were significantly correlated with SNR were removed from the feature set without correcting for multiple comparisons (*p*_spin_ < 0.05; 10 000 spatial autocorrelation-preserving permutation tests [70, 71]). Note that this is a more conservative feature selection procedure compared to conventional multiple comparisons correction, because fewer features would be removed if correction for multiple comparisons was applied. PCA was applied to the remaining set of features (Fig. S3b). The principal component of the retained 3 819 features (i.e., PC1 - feature subset) explained 31.6% of the variance and was significantly correlated with the original PC1 of the full set of features (*r*_s_ = 0.93, *p*_spin_ = 0.0001), reflecting similar spatial pattern as the original analysis. (2) SNR was regressed out from the full set of time-series features using linear regression analysis. PCA was then applied to the resulting feature residuals (Fig. S3c). The principal component of SNR-regressed features (i.e., PC1 - SNR regressed) explained 41.4% of the variance and reflected the same spatial pattern as the original analysis (*r*_s_ = 0.70, *p*_spin_ = 0.0004). Moreover, we assessed the effects of environmental and instrumental noise on the findings, where we applied principal component analysis to the `hctsa` features obtained from pre-processed empty-room MEG recordings [23] (see *Methods* for more details). PCA weights of the time-series features of the empty-room MEG recordings were aligned with the PCA weights of the time-series features of the resting-state MEG recordings using the Procrustes method ([89]; https://github.com/satra/mapalign). The principal component of neurophysiological dynamics was then compared with the principal component of time-series features obtained from empty-room recordings, where no significant associations were identified (Fig. S4; *r*_s_ = −0.17, *p*_spin_ = 0.69). These analyses demonstrate that the timeseries features captured by the dominant axis of variation in neurophysiological dynamics are independent from measures of MEG signal-to-noise ratio.

Finally, to ensure that the findings are independent from the parcellation resolution, we repeated the analyses using a higher resolution parcellation (Schaefer-400 atlas with 400 cortical regions [67]). The results were consistent with the original analysis (Fig. S5 and Fig. S6). The first principal component (PC1) accounted for 48.6% of the variance and displayed a similar spatial organization as the one originally obtained for the Schaefer-100 atlas (Fig. S5a). As before, the top loading time-series features were mainly related to the characteristics of the power spectral density (Fig. S5b,c). The full list of features, their loadings and *p*-values are available in the online Supplementary File S3. Moreover, PLS analysis identified a single significant latent variable (*p*_spin_ = 0.0083) that accounted for 75.7% of the covariance (Fig. S6a). Micro-architecture and timeseries feature scores displayed similar spatial patterns to the ones obtained for the Schaefer-100 atlas (Fig. S6b). The corresponding feature loadings were also consistent with the original findings (micro-architectural loadings in Fig. S6c and time-series feature loadings in the online Supplementary File S4.)

## DISCUSSION

In the present study, we use time-series phenotyping analysis to comprehensively chart the dynamic fingerprint of neurophysiological activity from the resting-state MEG signal. We then map the resulting dynamical atlas to a multimodal micro-architectural atlas to identify the neurophysiological signatures of cortical microarchitecture. We demonstrate that cortical variation in neurophysiological time-series properties mainly reflects power spectral density and is closely associated with intrinsic timescale and self-correlation structure of the signal. Moreover, the spatial organization of neurophysiological dynamics follows gradients of micro-architecture, such as neurotransmitter receptor and transporters, gene expression and T1w/T2w ratios, and reflects cortical metabolic demands.

Numerous studies have previously investigated neural oscillations and their relationship with neural communication patterns in the brain [8, 10, 11, 90]. Previous reports also suggest that neural oscillations influence behaviour and cognition [90–94] and are involved in multiple neurological diseases and disorders [93, 95]. Neural oscillations manifest as the variations of power amplitude of neurophysiological signal in the frequency domain [10, 21, 96, 97]. Power spectral characteristics of the neurophysiological signal, such as mean power amplitude in canonical frequency bands, have previously been used to investigate the underlying mechanisms of large-scale brain activity and to better understand the individual differences in brain function [22, 23, 31, 94, 98, 99]. Other time-series properties that are related to the power spectral density have also been used to study neural dynamics, including measures of intrinsic timescale and self-affinity or self-similarity of the signal (e.g., autocorrelation and fluctuation analysis) [25, 30, 77, 78, 100–102].

Applying a data-driven time-series feature extraction analysis, we find that the topographic organization of neurophysiological time-series signature follows a sensory-fugal axis, separating somatomotor, occipital and parietal cortices from anterior temporal, orbitofrontal and ventromedial cortices. This dynamic fingerprint of neurophysiological activity is mainly characterized by linear correlation structure of MEG signal captured by `hctsa` time-series features. The linear correlation structure manifests in both power spectral properties and the autocorrelation function. This dominant spatial variation of time-series features also resembles the spatial distribution of intrinsic timescale, another measure related to the characteristics of power spectral density [28, 30, 33]. Altogether, while the findings highlight under-represented time-series features, they emphasize the importance of conventional methods in characterizing neurophysiological activity and the key role of linear correlation structure in MEG dynamics.

Earlier reports found that regional neural dynamics, including measures of power spectrum and intrinsic timescale, reflect the underlying circuit properties and cortical micro-architecture [25, 28, 30]. The relationship between neural dynamics and cortical micro-architecture is often examined using a single, or a few microstructural features. Recent advances in data collection and integration and the increasing number of data sharing initiatives have provided a unique opportunity to comprehensively study cortical circuit properties and microarchitecture using a wide range of multimodal datasets [43, 44, 47, 56, 57, 65, 103]. Here we use such datasets and compile multiple micro-architectural maps, including measures of microstructure, metabolism, cortical expansion, receptors and transporters, layer thickness and cell type-specific gene expressions, to chart the multivariate associations between neurophysiological dynamics and cortical micro-architecture.

Our findings build on previous reports by showing that neurophysiological dynamics follow the underlying cytoarchitectonic and microstructural gradients. In particular, our findings confirm that MEG intrinsic dynamics are associated with the heterogeneous distribution of gene expression and myelination [30, 85] and neurotransmitter receptors and transporters [47]. In addition, we link the dynamic signature of ongoing neurophysiological activity with multiple metabolic attributes [62, 104]; for instance, we find that regions with greater oxygen and glucose metabolism tend to display lower temporal autocorrelation and therefore more variable moment-to-moment intrinsic activity. This is consistent with previously reported high metabolic rates of oxygen and glucose consumption in the sensory cortex [61]. We also find a prominent association with cell type-specific gene expression of oligodendrocyte precursors (opc), potentially reflecting the contribution of these cells to myelin generation by giving rise to myelinating oligodendrocytes during development [80–84] and to myelin regulation and metabolic support of myelinated axons in the adult neural circuits [83, 84, 105]. Finally, we find that the dominant dynamic signature of neural activity covaries with the granular cortical layer IV, consistent with the idea that layer IV receives prominent subcortical (including thalamic) feedforward projections [106, 107]. Collectively, our findings build on the emerging literature on how heterogeneous micro-architectural properties along with macroscale network embedding (e.g., cortico-cortical connectivity and subcortical projections) jointly shape regional neural dynamics [38–40, 108–111].

The present findings must be interpreted with respect to several methodological considerations. First, we used MEG data from a subset of individuals with no familial relationships from the HCP dataset. Although all the presented analyses are performed using the group-level data, future work with larger sample sizes can provide more generalizable outcomes [112, 113]. Larger sample sizes will also help go beyond associative analysis and allow for predictive analysis of neural dynamics and micro-architecture in unseen datasets. Second, MEG is susceptible to low SNR and has variable sensitivity to neural activity from different regions (i.e., sources). Thus, electrophysiological recordings with higher spatial resolution, such as intracranial electroencephalography (iEEG and ECoG), may provide more precise measures of neural dynamics that can be examined with respect to cortical micro-architecture. Finally, despite the fact that we attempt to use a near-comprehensive list of time-series properties and multiple micro-architectural features, neither the time-series features nor the micro-architectural maps are exhaustive sets of measures. Moreover, micro-architectural features are group-average maps that are compiled from different datasets. Multimodal datasets from the same individuals are required to perform individual-level comparisons between the dynamical and micro-architectural atlases.

Altogether, using a data-driven approach, the present findings show that neurophysiological signatures of cortical micro-architecture are hierarchically organized across the cortex, reflecting the underlying circuit properties. These findings highlight the importance of conventional measures for studying the characteristics of neurophysiological activity, while also identifying less-commonly used time-series features that covary with cortical micro-architecture. Collectively, this work opens new avenues for studying the anatomical basis of neurophysiological activity.

## METHODS

### Dataset: Human Connectome Project (HCP)

Resting state magnetoencephalography (MEG) data from a sample of healthy young adults (*n* = 33; age range 22–35 years) with no familial relationships were obtained from Human Connectome Project (HCP; S900 release [65]). The data includes resting state scans of approximately 6 minutes long (sampling rate = 2034.5 Hz; anti-aliasing low-pass filter at 400 Hz) and empty-room recordings for all participants. 3T structural magnetic resonance imaging (MRI) data and MEG anatomical data (i.e., cortical sheet with 8 004 vertices and transformation matrix required for co-registration of MEG sensors and MRI scans) of all participants were also obtained for MEG pre-processing.

### Resting state magnetoencephalography (MEG)

Resting state MEG data was analyzed using Brainstorm software, which is documented and freely available for download online under the GNU general public license ([66]; http://neuroimage.usc.edu/brainstorm). For each individual, MEG sensor recordings were registered to their structural MRI scan using the anatomical transformation matrix provided by HCP for co-registration, following the procedure described in Brainstorm online tutorials for the HCP dataset (https://neuroimage.usc.edu/brainstorm/Tutorials/HCP-MEG). The data were downsampled to 1/4 of the original sampling rate (i.e 509 Hz) to facilitate processing. The pre-processing was performed by applying notch filters at 60, 120, 180, 240, and 300 Hz, and was followed by a high-pass filter at 0.3 Hz to remove slow-wave and DC-offset artifacts. Bad channels were marked based on the information obtained through the data management platform of HCP (ConnectomeDB; https://db.humanconnectome.org/). The artifacts (including heartbeats, eye blinks, saccades, muscle movements, and noisy segments) were then removed from the recordings using automatic procedures as proposed by Brainstorm. More specifically, electrocardiogram (ECG) and electrooculogram (EOG) recordings were used to detect heartbeats and blinks, respectively. We then used Signal-Space Projections (SSP) to automatically remove the detected artifacts. We also used SSP to remove saccades and muscle activity as low-frequency (1-7 Hz) and high-frequency (40-240 Hz) components, respectively.

The pre-processed sensor-level data was then used to obtain a source estimation on HCP’s fsLR4k cortical surface for each participant (i.e., 8 004 vertices). Head models were computed using overlapping spheres and the data and noise covariance matrices were estimated from the resting state MEG and noise recordings. Linearly constrained minimum variance (LCMV) beamforming from Brainstorm was then used to obtain the source activity for each participant. We performed data covariance regularization to avoid the instability of data covariance matrix inversion due to the smallest eigenvalues of its eigen-spectrum. Data covariance regularization was performed using the “median eigenvalue” method from Brainstorm [66], such that the eigenvalues of the eigenspectrum of data covariance matrix that were smaller than the median eigenvalue were replaced with the median eigenvalue itself. The estimated source variance was also normalized by the noise covariance matrix to reduce the effect of variable source depth. Source orientations were constrained to be normal to the cortical surface at each of the 8 004 vertex locations on the fsLR4k surface. Source-level time-series were parcellated into 100 regions using the Schaefer-100 atlas [67] for each participant, such that a given parcel’s time series was estimated as the first principal component of its constituting sources’ time series. Finally, we estimated source-level signal-to-noise ratio (SNR) as follows [87, 88]:

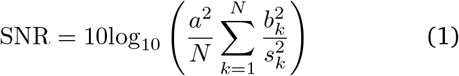

where *a* is the source amplitude (i.e., typical strength of a dipole, which is 10 nAm [114]), *N* is the number of sensors, *b_k_* is the signal at sensor *k* estimated by the forward model for a source with unit amplitude, and 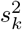 is the noise variance at sensor *k*. Group-average source-level SNR was parcellated using the Schaefer-100 atlas.

To estimate a measure of environmental and instrumental noise, empty-room MEG recordings of all individuals were obtained from HCP and were pre-processed using an identical procedure to the resting-state recordings. The pre-processed source-level time-series obtained from empty-room recordings were parcellated and subjected to time-series feature extraction analysis to estimate time-series features from noise data for each participant (see “Time-series feature extraction using `hctsa`”).

### Power spectral analysis

Welch’s method was used to estimate power spectrum density (PSD) from the source-level time-series for each individual, using overlapping windows of length 4 seconds with 50% overlap. Average power at each frequency band was then calculated for each vertex (i.e., source). Source-level power data were parcellated into 100 regions using the Schaefer-100 atlas [67] at six canonical electrophysiological bands (i.e., delta (*δ*: 2-4 Hz), theta (*θ*: 5-7 Hz), alpha (*α*: 8-12 Hz), beta (*β*: 15-29 Hz), low gamma (lo-*γ*: 30-59 Hz), and high gamma (hi-*γ*: 60-90Hz)). We contributed the group-average vertex-level power maps on the fsLR4k surface to the publicly available neuromaps toolbox [43].

### Intrinsic timescale

The regional intrinsic timescale was estimated using spectral parameterization with the FOOOF (fitting oscillations & one over f) toolbox [21]. Specifically, the source-level power spectral density were used to extract the neural timescale at each vertex and for each individual using the procedure described by Gao et al. [30]. The FOOOF algorithm decomposes the power spectra into periodic (oscillatory) and aperiodic (1/*f*-like) components by fitting the power spectral density in the log-log space [21] (Fig. S1). The algorithm identifies the oscillatory peaks (periodic component), the “knee parameter”*k* that controls for the bend in the aperiodic component and the aperiodic “exponent”*χ* [21, 30]. The knee parameter *k* is then used to calculate the “knee frequency” as *f_k_* = *k*^1/*χ*^, which is the frequency where a knee or a bend occurs in the power spectrum density [30]. Finally, the intrinsic timescale *τ* is estimated as [30]:

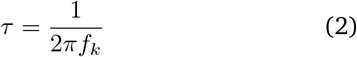

We used the FOOOF algorithm to fit the power spectral density with “knee” aperiodic mode over the frequency range of 1-60 Hz. Note that since the first notch filter was applied at 60 Hz during the pre-processing analysis, we did not fit the model above 60 Hz. Following the guidelines from the FOOOF algorithm and Donoghue et al. [21], the rest of the parameters were defined as: peak width limits (peak_width_limits) = 1-6 Hz; maximum number of peaks (max_n_peaks) = 6; minimum peak height (min_peak_height) = 0.1; and peak threshold (peak_threshold) = 2. Intrinsic timescale *τ* was estimated at each vertex for each individual and was parcellated using the Schaefer-100 atlas [67]. We contributed the group-average vertex-level intrinsic timescale map on the fsLR4k surface to the publicly available neuromaps toolbox [43].

In addition to the aperiodic component used to calculate the intrinsic timescale, the FOOOF spectral parameterization algorithm also provides the extracted peak parameters of the periodic component at each vertex for each participant. We used the oscillatory peak parameters to estimate band-limited power maps that were adjusted for the aperiodic component as opposed to the total power maps estimated above [21] (see *Power spectral analysis*). We defined the power band limits as delta (2-4 Hz), theta (5-7 Hz), alpha (8-14 Hz), and beta (15-30 Hz), based on the distribution of peak center frequencies across all vertices and participants (Fig. S1b). Given the lack of clear oscillatory peaks in high frequencies (above 40 Hz), the FOOOF algorithm struggles with detecting consistent peaks in gamma frequencies and above [21, 22]. Thus, we did not analyze band-limited power in gamma frequencies using spectral parameterization. For each of the 4 predefined power bands, we estimated an “oscillation score” following the procedure described by Donoghue et al. [21]. Specifically, for each participant and frequency band, we identified the extracted peak at each vertex. If more than one peak was detected at a given vertex, the peak with maximum power was selected. The average peak power was then calculated at each vertex and frequency band across participants. The group-average peak power map was then normalized for each frequency band, such that the average power at each vertex was divided by the maximum average power across all vertices. Separately, we calculated a vertex-level probability map for each frequency band as the percentage of participants with at least one detected peak at a given vertex at that frequency band. Finally, the bandlimited “oscillation score” maps were obtained by multiplying the normalized group-average power maps with their corresponding probability maps for each frequency band. The oscillation score maps were parcellated using the Schaefer-100 atlas [67] (Fig. S1a).

### Time-series feature extraction using hctsa

We used the highly comparative time-series analysis toolbox, `hctsa` [35, 36], to perform a massive feature extraction of the pre-processed time-series for each brain region for each participant. The `hctsa` package extracted over 7 000 local time-series features using a wide range of time-series analysis methods [35, 36]. The extracted features include, but are not limited to, measures of data distribution, temporal dependency and correlation properties, entropy and variability, parameters of time-series model fit, and nonlinear properties of a given time-series [35, 37].

The `hctsa` feature extraction analysis was performed on the parcellated MEG time-series for each participant. Given that applying `hctsa` on the full time-series is computationally expensive, we used 80 seconds of data for feature extraction after dropping the first 30 seconds. Previous reports suggest that relatively short segments of about 30 to 120 seconds of resting-state data are sufficient to estimate robust properties of intrinsic brain activity [22]. Nevertheless, to ensure that we can robustly estimate time-series features from 80 seconds of data, we calculated a subset of `hctsa` features using the catch-22 toolbox [115] on subsequent segments of time-series with varying length for each participant. Specifically, we extracted time-series features from short segments of data ranging from 5 to 125 seconds in increments of 5 seconds. To identify the time-series length required to estimate robust and stable features, we calculated the Pearson correlation coefficient between features of two subsequent segments (e.g., features estimated from 10 and 5 seconds of data). The correlation coefficient between the estimated features started to stabilize at time-series segments of around 30 seconds, consistent with previous reports [22] (Fig. S7). Following the feature extraction procedure from time-series segments of 80 seconds, the outputs of the operations that produced errors were removed and the remaining features (6 880 features) were normalized across nodes using an outlier-robust sigmoidal transform for each participant separately. A group-average region × feature matrix was generated from the normalized individual-level features. We also applied `hctsa` analysis to the parcel-lated empty-room recordings (80 seconds) to estimate time-series features from noise data using an identical procedure to resting-state data, identifying 6 148 features per region per participant. The time-series features were normalized across brain regions for each participant. A group-average empty-room feature set was obtained and used for further analysis.

### Micro-architectural features from neuromaps

We used the neuromaps toolbox (https://github.com/netneurolab/neuromaps) [43] to obtain micro-architectural and neurotransmitter receptor and transporter maps in the maps’ native spaces. Details about all maps and their data sources are available in [43]. Briefly, all data that were originally available in any surface space were transformed to the fsLR32k surface space using linear interpolation to resample data and were parcellated into 100 cortical regions using the Schaefer-100 atlas in fsLR32k space [67]. All volumetric data were retained in their native MNI152 volumetric space and were parcellated into 100 cortical regions using the volumetric Schaefer atlas in MNI152 space [67]. Micro-architectural maps included T1w/T2w as a proxy measure of cortical myelin [116], cortical thickness [116], principal component of gene expression [44, 68], principal component of neurotransmitter receptors and transporters [47], synapse density (using [^11^*C*]UCB-J PET tracer that binds to the synaptic vesicle glycoprotein 2A (SV2A)) [55, 117–128], metabolism (i.e., cerebral blood flow and volume, oxygen and glucose metabolism, glycolytic index) [61], evolutionary and developmental expansion [63], allometric scaling from Philadelphia Neurodevelopmental Cohort (PNC) and National Institutes of Health (NIH) [64]. Neurotransmitter maps included 18 different neurotransmitter receptors and transporters across 9 different neurotransmitter systems, namely serotonin (5-HT1a, 5-HT1b, 5-HT2a, 5-HT4, 5-HT6, 5-HTT), histamine (H3), dopamine (D1, D2, DAT), norepinephrine (NET), acetylcholine (*α*4*β*2, M1, VAChT), cannabinoid (CB1), opioid (MOR), glutamate (mGluR5), and GABA (GABAa/bz) [47].

### BigBrain histological data

Layer thickness data for the 6 cortical layers (I-VI) were obtained from the BigBrain atlas, which is a volumetric, high-resolution (20×20×20*μ*m) histological atlas of a post-mortem human brain (65-year-old male) [56–58]. In the BigBrain atlas, sections of the post mortem brain are stained for cell bodies using Merker staining technique [129]. These sections are then imaged and used to reconstruct a volumetric histological atlas of the human brain that reflects neuronal density and soma size and captures the regional differentiation of cytoarchitecture [56–58, 103, 130]. The approximate cortical layer thickness data obtained from the BigBrainWarp toolbox [57], were originally generated using a convolutional neural network that automatically segments the cortical layers from the pial to white surfaces [58]. Full description of how the cortical layer thickness was approximated is available elsewhere [58]. The cortical layer thickness data for the 6 cortical layers were obtained on the *fsaverage* surface (164k vertices) from the BigBrainWarp toolbox [57] and were parcellated into 100 cortical regions using the Schaefer-100 atlas [67].

### Cell type-specific gene expression

Regional microarray expression data were obtained from 6 post-mortem brains (1 female, ages 24–57, 42.5 ± 13.4) provided by the Allen Human Brain Atlas (AHBA, https://human.brain-map.org; [44]). Data were processed with the abagen toolbox (version 0.1.3-doc; https://github.com/rmarkello/abagen; [68]) using the Schaefer-100 volumetric atlas in MNI space [67].

First, microarray probes were reannotated using data provided by [131]; probes not matched to a valid Entrez ID were discarded. Next, probes were filtered based on their expression intensity relative to background noise [132], such that probes with intensity less than the background in ≥ 50.00% of samples across donors were discarded. When multiple probes indexed the expression of the same gene, we selected and used the probe with the most consistent pattern of regional variation across donors (i.e., differential stability; [133]), calculated with:

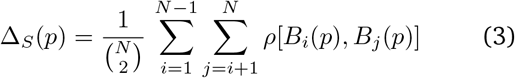

where *ρ* is Spearman’s rank correlation of the expression of a single probe, *p*, across regions in two donors *B_i_* and *B_j_*, and N is the total number of donors. Here, regions correspond to the structural designations provided in the ontology from the AHBA.

The MNI coordinates of tissue samples were updated to those generated via non-linear registration using the Advanced Normalization Tools (ANTs; https://github.com/chrisfilo/alleninf). To increase spatial coverage, tissue samples were mirrored bilaterally across the left and right hemispheres [134]. Samples were assigned to brain regions in the provided atlas if their MNI coordinates were within 2 mm of a given parcel. If a brain region was not assigned a tissue sample based on the above procedure, every voxel in the region was mapped to the nearest tissue sample from the donor in order to generate a dense, interpolated expression map. The average of these expression values was taken across all voxels in the region, weighted by the distance between each voxel and the sample mapped to it, in order to obtain an estimate of the parcellated expression values for the missing region. All tissue samples not assigned to a brain region in the provided atlas were discarded.

Inter-subject variation was addressed by normalizing tissue sample expression values across genes using a robust sigmoid function [35]:

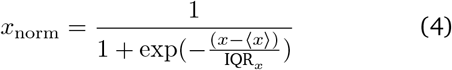

where 〈*x*〉 is the median and IQR_*x*_ is the normalized interquartile range of the expression of a single tissue sample across genes. Normalized expression values were then rescaled to the unit interval:

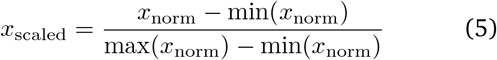

Gene expression values were then normalized across tissue samples using an identical procedure. Samples assigned to the same brain region were averaged separately for each donor and then across donors, yielding a regional expression matrix of 15,633 genes.

Finally, cell type-specific gene expression maps were calculated using gene sets identified by a cell type deconvolution analysis [59, 60, 76]. Detailed description of the analysis is available at [59]. Briefly, cell-specific gene sets were compiled across 5 single-cell and single-nucleus RNA sequencing studies of adult human post-mortem cortical samples [135–140]. Gene expression maps of the compiled study-specific cell types were obtained from AHBA. Unsupervised hierarchical clustering analysis was used to identify 7 canonical cell classes that included astrocytes (astro), endothelial cells (endo), microglia (micro), excitatory neurons (neuron-ex), inhibitory neurons (neuron-in), oligodendrocytes (oligo) and oligodendrocyte precursors (opc) [59]. We used the resulting gene sets to obtain average cell type-specific expression maps for each of these 7 cell classes from the regional expression matrix of 15 633 genes.

### Partial Least Squares (PLS)

Partial least squares (PLS) analysis was used to investigate the relationship between resting-state MEG timeseries features and micro-architecture maps. PLS is a multivariate statistical technique that identifies mutually orthogonal, weighted linear combinations of the original variables in the two datasets that maximally covary with each other, namely the latent variables [74, 75]. In the present analysis, one dataset is the `hctsa` feature matrix (i.e., **X**_*n*×*t*_) with *n* = 100 rows as brain regions and *t =* 6880 columns as time-series features. The other dataset is the compiled set of micro-architectural maps (i.e., **Y**_*n*×*m*_) with *n* = 100 rows (brain regions) and *m* = 45 columns (micro-architecture maps). To identify the latent variables, both data matrices were normalized column-wise (i.e., *z*-scored) and a singular value decomposition was applied to the correlation matrix **R** = **X′ Y** as follows:

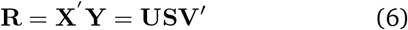

where **U**_*t*×*m*_ and **V**_*m*×*m*_ are orthonormal matrices of left and right singular vectors and **S**_*m*×*m*_ is the diagonal matrix of singular values. Each column of **U** and **V** matrices corresponds to a latent variable. Each element of the diagonal of **S** is the corresponding singular value. The singular values are proportional to the covariance explained by latent variable and can be used to calculate effect sizes as 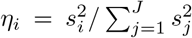 where *η_i_* is the effect size for the i-th latent variable (LV_*i*_), *s_i_* is the corresponding singular value, and *J* is the total number of singular values (here *J* = *m*). The left and right singular vectors **U** and **V** demonstrate the extent to which the time-series features and micro-architectural maps contribute to latent variables, respectively. Time-series features with positive weights covary with micro-architectural maps with positive weights, while negatively weighted time-series features and micro-architectural maps covary together. Singular vectors can be used to estimate brain scores that demonstrate the extent to which each brain region expresses the weighted patterns identified by latent variables. Brain scores for time-series features and micro-architectural maps are calculated by projecting the original data onto the PLS-derived weights (i.e., **U** and **V**):

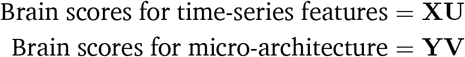

Loadings for time-series features and micro-architectural maps are then computed as the Pearson correlation coefficient between the original data matrices and their corresponding brain scores. For example, time-series feature loadings are the correlation coefficients between the original `hctsa` time-series feature vectors and PLS-derived brain scores for time-series features.

The statistical significance of latent variables was assessed using 10 000 permutation tests, where the original data was randomized using spatial autocorrelation-preserving nulls (see “Null model” for more details). The PLS analysis was repeated for each permutation, resulting in a null distribution of singular values. The significance of the original singular values were then assessed against the permuted null distributions (Fig. 4a). The reliability of PLS loadings was estimated using bootstrap resampling [141], where rows of the original data matrices **X** and **Y** are randomly resampled with replacement 10 000 times. The PLS analysis was then repeated for each resampled data, generating a sampling distribution for each time-series feature and micro-architectural map (i.e., generating 10 000 bootstrap-resampled loadings). The bootstrap-resampled loading distributions are then used to estimate 95% confidence intervals for loadings (e.g., see Fig. 4d).

Finally, given that PLS-derived brain scores are by design highly correlated, we used a distance-dependent cross-validation analysis to assess the out-of-sample correlations between brain scores [76]. Specifically, 75% of the closest brain regions in Euclidean distance to a random “seed” region were selected as training set, while the 25% remaining distant regions were selected as test set. We then re-ran the PLS analysis on the training set (i.e., 75% of regions) and used the resulting weights (i.e., singular values) to estimated brain scores for test set. The out-of-sample correlation was then calculated as the Spearman’s rank correlation coefficient between test set brain scores of time-series features and micro-architectural maps. We repeated this analysis 99 times, such that each time a random brain region was selected as the seed region, yielding distributions of training set brain scores correlations and test set (out-of-sample) correlations (Fig. 4b). Note that 99 is the maximum number of train-test splits here given that brain maps consist of 100 regions.

### Null model

To make inferences about the topographic correlations between any two brain maps, we implement a null model that systematically disrupts the relationship between two topographic maps but preserves their spatial autocorrelation [70, 71, 142]. We used the Schaefer-100 atlas in the HCP’s fsLR32k grayordinate space [65, 67]. The spherical projection of the fsLR32k surface was used to define spatial coordinates for each parcel by selecting the vertex closest to the center-of-mass of each parcel [143–145]. The resulting spatial coordinates were used to generate null models by applying randomly-sampled rotations and reassigning node values based on the closest resulting parcel (10 000 repetitions). The rotation was applied to one hemisphere and then mirrored to the other hemisphere. Where appropriate, the results were corrected for multiple comparisons by controlling the false discovery-rate (FDR correction [146]).

## Code and data availability

Code used to conduct the reported analyses is available on GitHub (https://github.com/netneurolab/shafiei_megdynamics). Data used in this study were obtained from the Human Connectome Project (HCP) database (available at https://db.humanconnectome.org/).

## ACKNOWLEDGMENTS

We thank Justine Hansen, Estefany Suarez, Filip Milisav, Andrea Luppi, Vincent Bazinet, Zhen-Qi Liu for their comments on the manuscript. BM acknowledges support from the Natural Sciences and Engineering Research Council of Canada (NSERC), Canadian Institutes of Health Research (CIHR), Brain Canada Foundation Future Leaders Fund, the Canada Research Chairs Program, the Michael J. Fox Foundation, and the Healthy Brains for Healthy Lives initiative. SB acknowledges support from the NIH (R01 EB026299), a Discovery grant from the Natural Science and Engineering Research Council of Canada (NSERC 436355-13), the CIHR Canada research Chair in Neural Dynamics of Brain Systems, the Brain Canada Foundation with support from Health Canada, and the Innovative Ideas program from the Canada First Research Excellence Fund, awarded to McGill University for the Healthy Brains for Healthy Lives initiative. BV acknowledges support from NIH National Institute of General Medical Sciences grant (R01GM134363). GS acknowledges support from the Natural Sciences and Engineering Research Council of Canada (NSERC) and the Fonds de recherche du Québec - Nature et Technologies (FRQNT).

---

**Figure S1.**
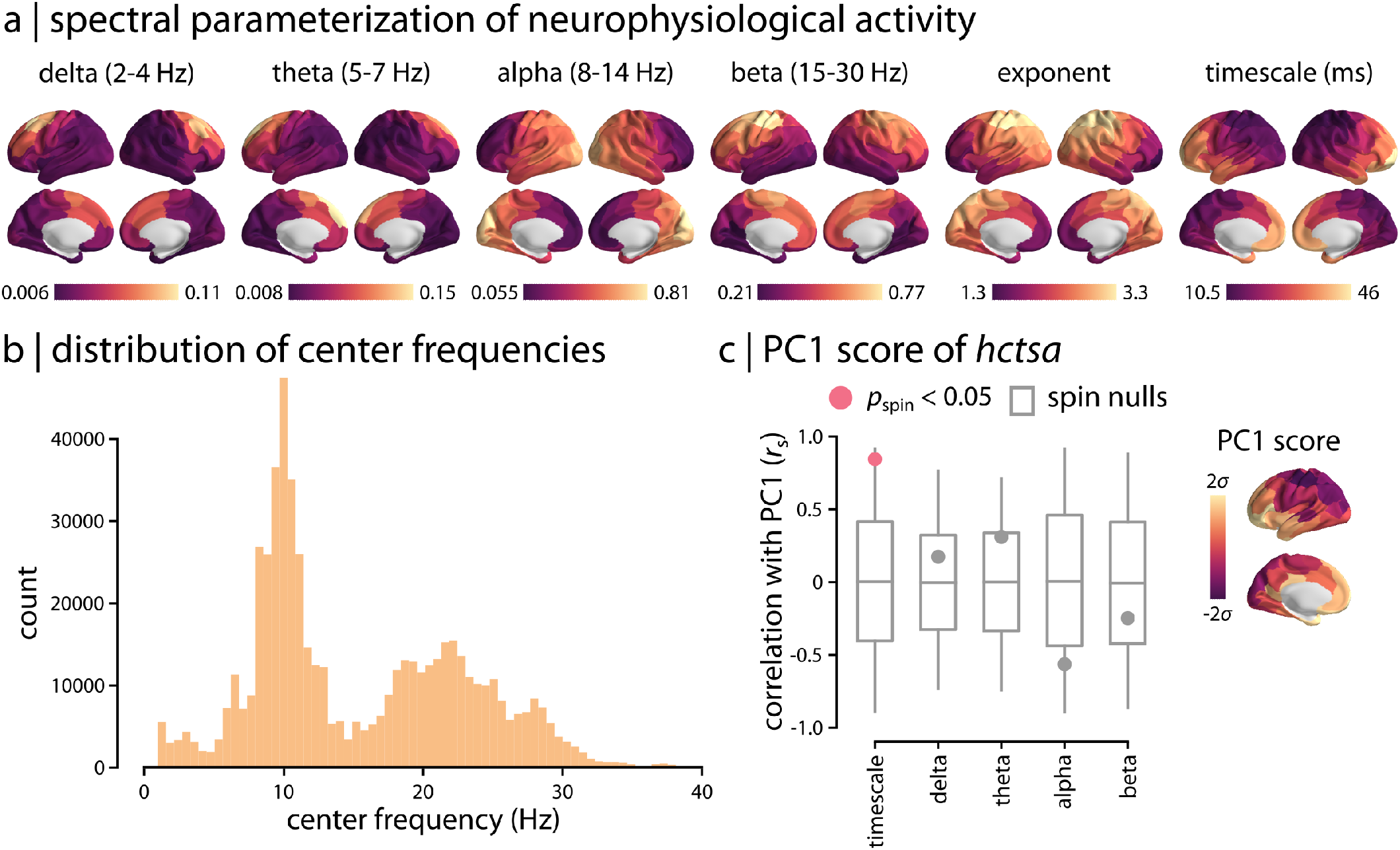
Spectral parameterization of neurophysiological activity. Spectral parameterization FOOOF toolbox was used to extract periodic and aperiodic components of the MEG power spectrum [21]. (a) The identified oscillatory peaks of the periodic component were used to estimate band-limited “oscillation score” maps at delta (2-4 Hz), theta (5-7 Hz), alpha (8-14 Hz), and beta (15-30 Hz) frequency bands. The oscillation scores reflect the average power at each region and frequency band, weighted by the probability of observing an oscillation peak at that region and band [21]. The aperiodic exponent and “knee” parameter (controls for the bend in the aperiodic component) were used to estimate intrinsic timescale (see *Methods* for details). Note that log-10 transformed intrinsic timescale map is depicted here for a more clear visualization. (b) Distribution of center frequencies of the identified periodic peaks are depicted across all vertices and participants. Visual inspection of the distribution shows clusters of peaks around the frequency bands shown in panel (a). (c) PC1 score map of `hctsa` time-series features was compared with the aperiodic-adjusted power maps and intrinsic timescale. Consistent with the results obtained with the total power maps at the canonical frequencies (Fig. 3), PC1 is significantly associated with the intrinsic timescale (FDR-corrected; 10 000 autocorrelation-preserving spin nulls). *r*_s_ denotes the Spearman’s rank correlation coefficient.

**Figure S2.**
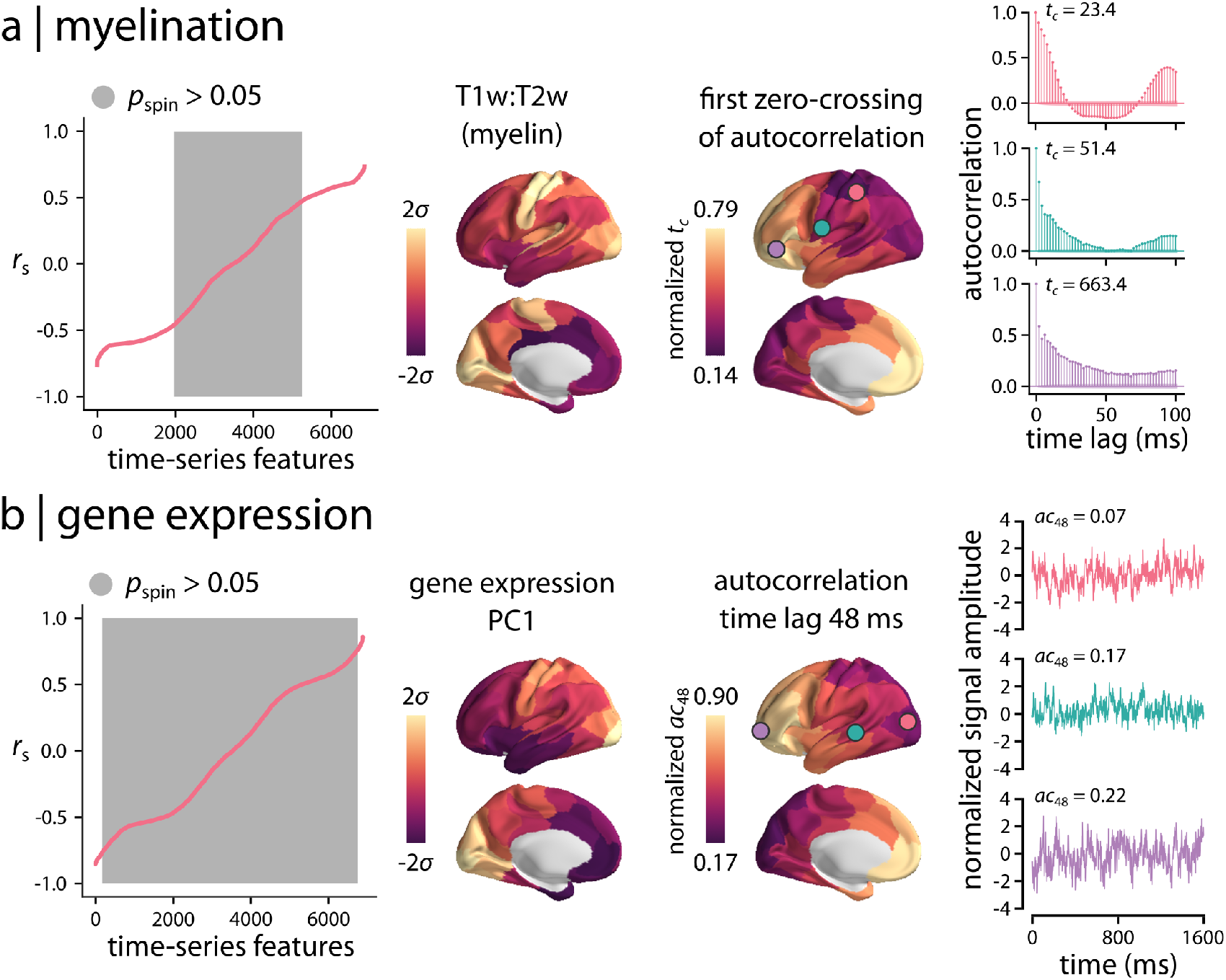
Univariate analysis of neurophysiological time-series features. Spearman’s rank correlation coefficients (r_s_) were used to investigate the univariate associations between `hctsa` time-series features of neurophysiological signal and two commonly-used micro-architectural maps: (a) T1w/Tw2 ratio as a proxy measure of myelination, and (b) principal component of gene expression. The resulting correlations were compared with null distributions of correlations obtained from 10 000 spatial autocorrelation-preserving nulls. Grey background indicates non-significant time-series features (FDR corrected). Examples of high loading time-series features are shown for each micro-architectural map. The group-average first zero-crossing time point of the autocorrelation function, *t_c_*, is shown for T1w/T2w ratio. The group-average linear autocorrelation at a time lag of 48 ms, *ac*_48_, is shown for principal component of gene expression. The autocorrelation function and short segments of raw time-series are also shown for a randomly selected participant at three different regions (circles on the brain surface: pink ≈ *5^th^* percentile, green ≈ 50^*th*^ percentile, purple ≈ 95^*th*^ percentile). Full lists of features, their correlation coefficients and *p*-values are available for T1w/T2w ratio and gene expression in the online Supplementary Files S5,6.

**Figure S3.**
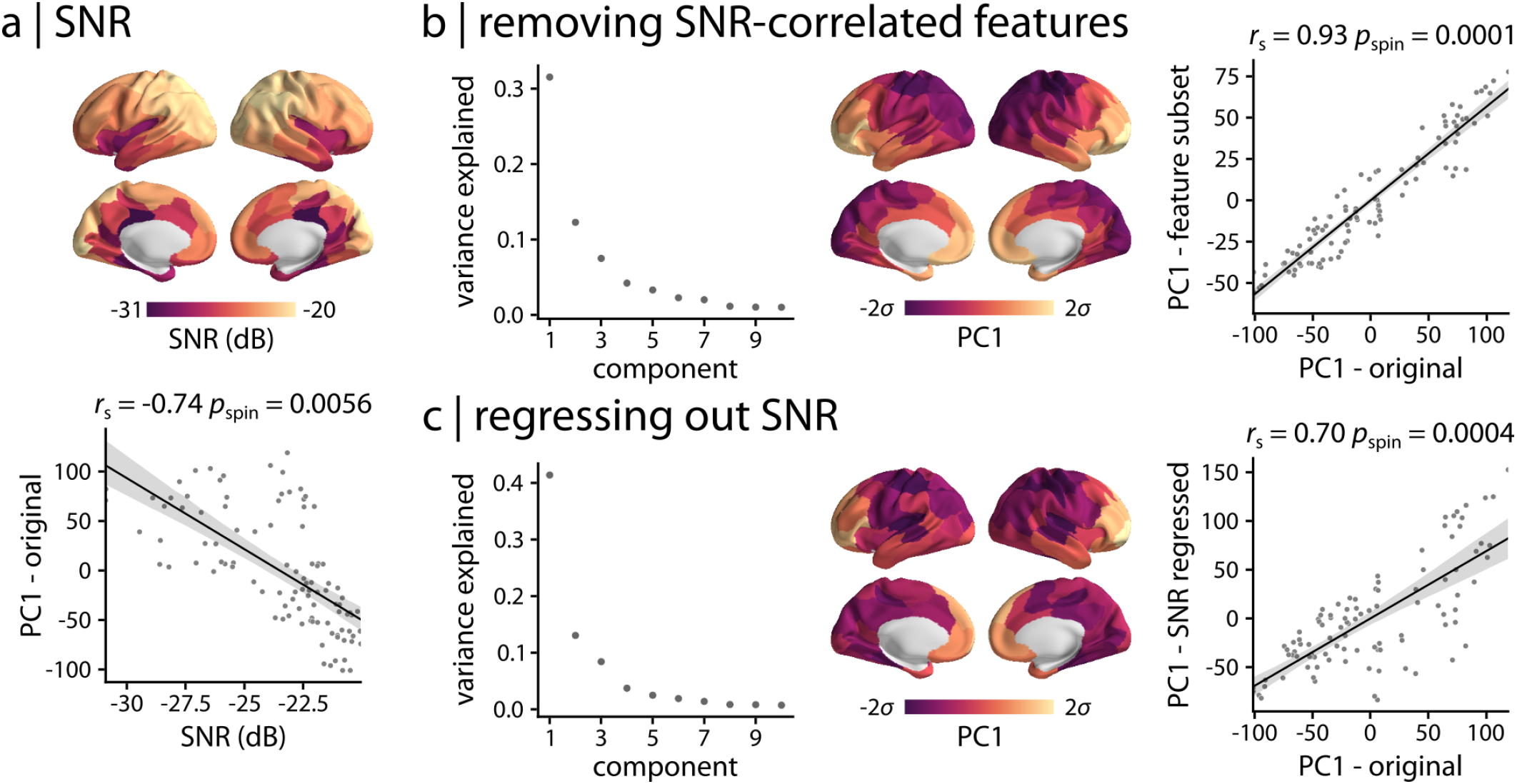
Signal-to-noise ratio (SNR) (a) Source-level MEG signal-to-noise ratio (SNR) was estimated. Parcellated, group-average SNR map is depicted across the cortex. MEG SNR was compared with the principal component of neurophysiological dynamics (PC1 - original). (b) SNR was compared with full set of neurophysiological time-series features (i.e., 6 880 features) using univariate correlations. Features that were significantly correlated with SNR were removed without correcting for multiple comparisons (*p*_spin_ < 0.05, 10 000 spatial autocorrelation-preserving permutation tests) and PCA was repeated using the remaining 3 819 features. The principal component of the retained feature subset (PC1 - feature subset) explained 31.6% of the variance and was significantly correlated with the original PC1 from the full set of features. (c) SNR was linearly regressed out from the full set of time-series features. PCA was applied to the feature residuals. The principal component of SNR-regressed time-series features (PC1 - SNR regressed) explained 41.4% of the variance and was significantly correlated with the original PC1. *r*_s_ denotes the Spearman’s rank correlation coefficient; linear regression lines are added to the scatter plots for visualization purposes only.

**Figure S4.**
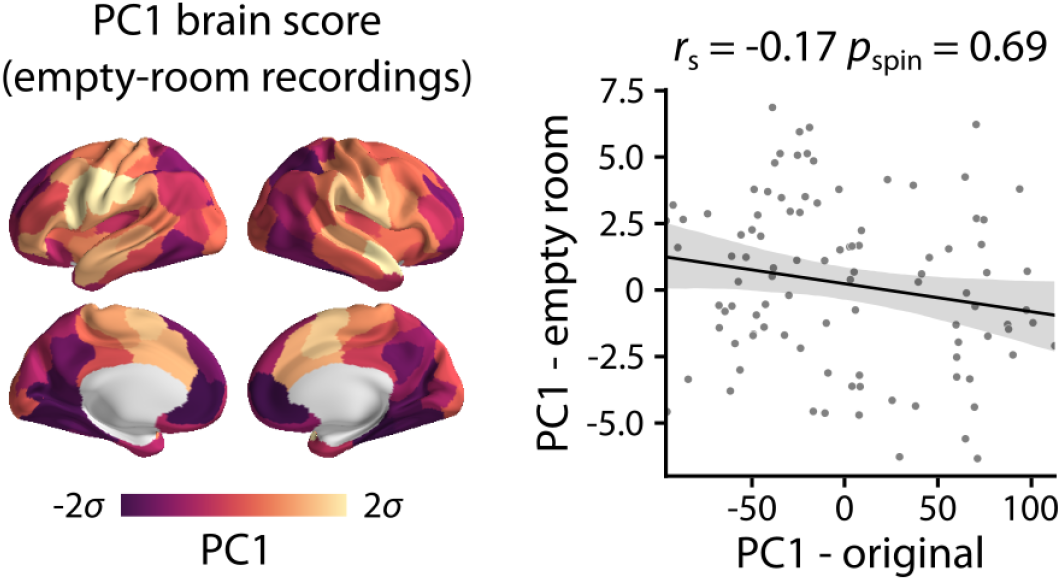
Empty-room recordings. PCA was applied to time-series features obtained from pre-processed empty-room MEG recordings. Following the Procrustes alignment of the resulting PCA weights with the PCA weights of the resting-state MEG recordings, the first principal components were compared between the two. The principal component of the empty-room timeseries features (PC1 - empty room; variance explained = 41%) was not significantly correlated with the original PC1. *r*_s_ denotes the Spearman’s rank correlation coefficient; linear regression line is added to the scatter plot for visualization purposes only.

**Figure S5.**
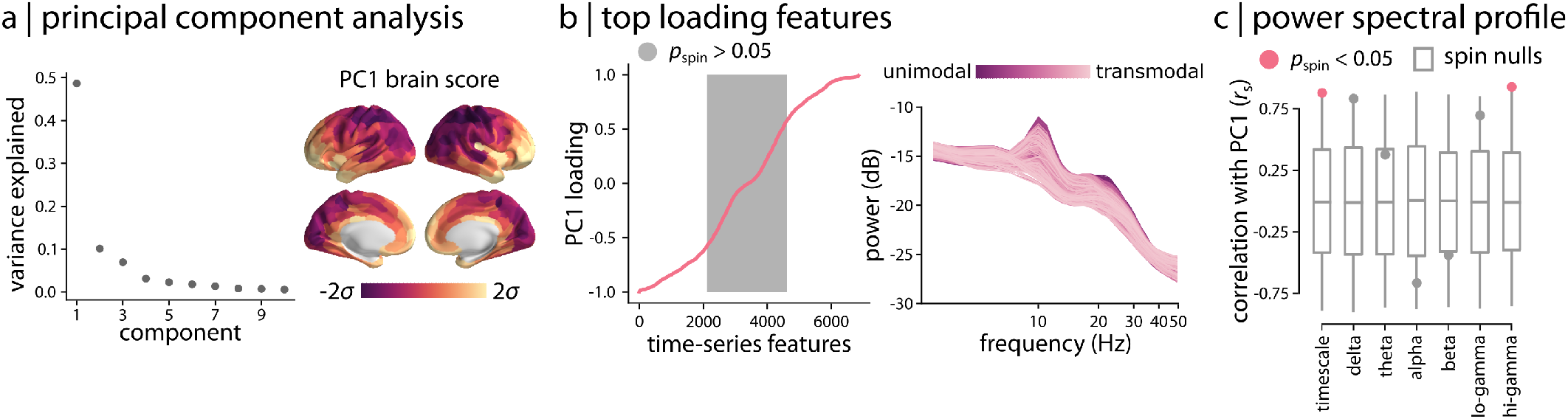
Topographic distribution of neurophysiological dynamics for Schaefer-400. (a) Principal component analysis (PCA) was applied to MEG time-series features at a higher resolution parcellation (i.e., Schaefer-400 atlas). The first principal component accounted for 48.6% of the variance. The spatial organization of features captured by PC1 is depicted across the cortex, displaying a consistent pattern with the original PCA results obtained for the Schaefer-100 atlas (Fig. 3). (b) Top loading features contributing to PC1 were identified using Pearson correlation coefficients between PC1 pattern and all time-series features. Grey background indicates non-significant features based on 10 000 spatial autocorrelation-preserving permutation tests (FDR corrected). Consistent with the original analysis, the top loading features were mainly related to power spectral density. Regional power spectral densities are depicted, where each line represents a brain region. Regions are coloured by their position in the putative unimodal–transmodal hierarchy [72]. The full list of features, their loadings and *p*-values are available in the online Supplementary File S3. (c) PC1 pattern was directly correlated with MEG power maps at 6 canonical frequency bands and intrinsic timescales. Consistent with the original findings, PC1 score was significantly correlated with hi-gamma power and intrinsic timescale (FDR-corrected). *r*_s_ denotes the Spearman’s rank correlation coefficient.

**Figure S6.**
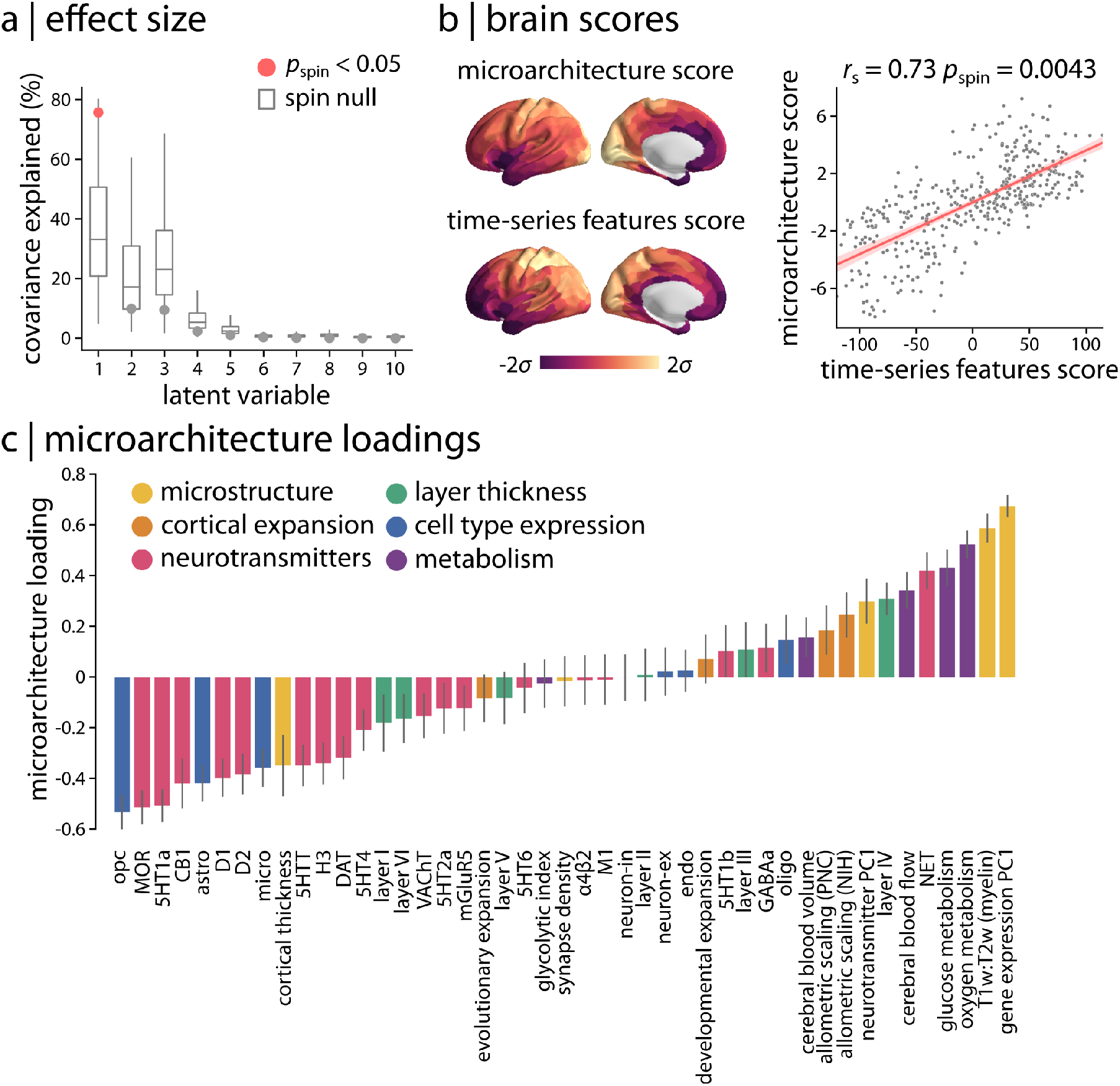
Partial least squares (PLS) analysis for Schaefer-400. PLS analysis was used to assess the multivariate relationship between micro-architectural and time-series features for the Schaefer-400 atlas. The results were consistent with the original findings for the Schaefer-100 atlas (Fig. 4). (a) PLS identified a single significant latent variable (*p*_spin_ = 0.0083, covariance explained = 75.7%). (b) Spatial patterns of micro-architecture and time-series features scores are depicted for the first latent variable. The two brain score maps were significantly correlated, demonstrating similar patterns to the ones obtained for Schaefer-100 atlas (Fig. 4b). (c) Micro-architectural feature loadings were also consistent with the original findings (Fig. 4d). Full list of time-series feature loadings are included in the online Supplementary File S4. Consistent with the original analysis, the top loading features were mainly related to the linear correlation structure of the signal. *r*_s_ denotes the Spearman’s rank correlation coefficient.

**Figure S7.**
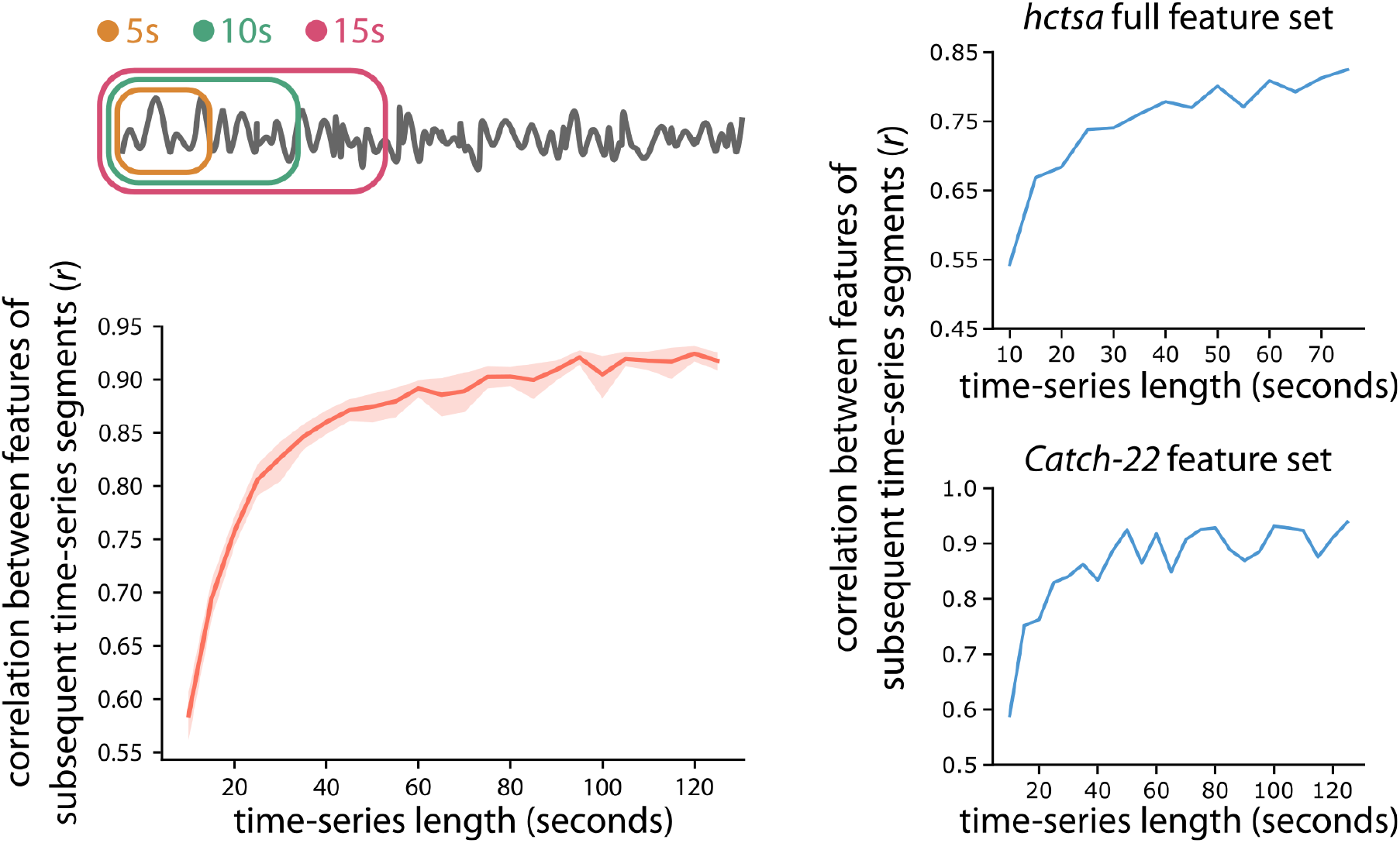
Stability of time-series features. To identify the time-series length required to robustly estimate the time-series features, we calculated a subset of `hctsa` features using the catch-22 toolbox [115] on subsequent segments of time-series with varying length for each participant. We extracted time-series features from short segments of data ranging from 5 to 125 seconds in increments of 5 seconds. To identify the optimal time-series length required to estimate robust and stable features, we calculated the Pearson correlation coefficient *r* between features of two subsequent segments (e.g., features estimated from 10 and 5 seconds of data). The group-average correlation coefficient between the estimated features started to stabilize at time-series segments of around 30 seconds, consistent with previous reports [22] (left). To compare the stability analysis of catch-22 features with full `hctsa` features, the correlation coefficients between subsequent segments of time-series are shown for a randomly selected participant (right).

## Notes

### Competing Interest Statement

The authors have declared no competing interest.

